# Landscape variation in defense traits along gradients of multiple resources and mammalian herbivory

**DOI:** 10.1101/2022.10.28.514290

**Authors:** Neha Mohanbabu, Michiel P Veldhuis, Dana Jung, Mark E Ritchie

## Abstract

Variation in defense traits likely depends on access to different resources and risk from herbivory. Plant defense theories have predicted both positive and negative associations between defense traits and access to resources, but relatively few studies have explored intraspecific variation in defense traits along multiple resource and mammalian herbivory risk gradients. We assessed relationships between herbivory intensity, multiple resources, and plant defense traits using a widely distributed tropical savanna herb, *Solanum incanum*. As independent measures of risk from large mammal herbivores are rare, we used a satellite-based vegetation index to predict herbivory intensity at the landscape scale. We found that the satellite-based estimate of herbivory intensity was positively associated with browser abundance and total soil P, but negatively associated with rainfall. Intraspecific defense traits too varied substantially across sites (n=43) but only variation in spine density was associated with herbivory intensity and plant resources, such that spine density was positively associated with both rainfall and soil P, but bimodally associated with herbivory intensity. Taken together, it suggests that defenses maybe favored either where resources for defense are abundant under low but still present risk (i.e, at high rainfall sites) or where resource-expensive plant tissue is at high risk (i.e, at high soil P sites). This hints at the possibility of a shift from a resource-associated (bottom-up) to an herbivory-associated (top-down) control of allocation to defenses along an environmental gradient. Additionally, the independent effect of soil P on a carbon-based defense, spine density, suggests potential for resources that are not components of defenses to also influence allocation to defense traits. Thus, our study provides evidence for the influence of multiple drivers, resources, and herbivory intensity, on anti-herbivore defenses and their shifting relative importance on allocation to defenses along an environmental gradient.

## Introduction

Plants invest in a variety of anti-herbivore defenses such as physical (Hanley et al. 2007, Barton 2016), chemical (Levin 1976, Bennett and Wallsgrove 1994) and mutualistic defenses (Heil and McKey 2003). Even within species, there is a significant amount of variation in the type and quantity of defense which may be driven by access to resources (Castillo et al. 2013, Abdala-Roberts et al. 2016, Lynn and Fridley 2019). The availability of resources to plants may influence the costs and benefits of allocating resources to defense traits rather than plant growth and reproduction. These cost-benefit relationships can be further modified by risks from the type of herbivores (insect vs mammal, generalist vs specialist) and intensity of herbivory. Therefore, any resource that influences plant and/or herbivore growth may indirectly affect resource allocation to defense traits. Although plants and herbivores have been shown to be simultaneously limited by multiple resources such as water/light, nitrogen (N), and phosphorus (P)(Elser et al. 2007, Harpole et al. 2011, Sperfeld et al. 2012), studies on variation in defense traits along multiple resource gradients are relatively rare. Given the growing evidence of the importance of resources other than carbon (C) and N to both plants and herbivores (Fay 2015, Kaspari et al. 2017, Prather et al. 2018, Borer et al. 2019), it is crucial to understand their impact on plant defense traits.

The type of defense traits and its association with resource gradients may vary depending on the identity of the resource. For example, the long-standing carbon-nutrient balance hypothesis (CNBH) (Bryant et al. 1983)(Fig 1, orange line) predicts that, irrespective of responses of herbivores to resources, increased availability of carbon (via greater light, water, or CO_2_) should promote carbon-based defenses, such as phenolics which can be toxic and/or lignin which increases leaf toughness and reduces palatability. In contrast, increased availability of N should promote N-based defenses such as alkaloids but result in a decline of C-based defenses. Studies exploring variation in chemical defenses along gradients of light, water, and N have found some support for the CNBH (Larsson et al. 1986, Hemming and Lindroth 1999, Hamilton et al. 2001, Gowda et al. 2003, Massey et al. 2005). Despite the long residence of the CNBH in the literature, few studies have addressed whether plant resources other than C and N would influence plant defense traits (De Long et al. 2016). For instance, resources such as phosphorus (P), sodium (Na) or calcium (Ca), have been shown to increase herbivore abundances or intensity due to their impact on plant quantity and/or quality (i.e. tissue stoichiometry)(Bishop et al. 2010, Joern et al. 2012, La Pierre and Smith 2016, Kaspari et al. 2017, Prather et al. 2018, Kaspari 2020). But whether these resources that may not be structural components of plant defenses can indirectly influence allocation to defense traits via their impacts on herbivore abundance/intensity remains elusive.

**Figure 1:**
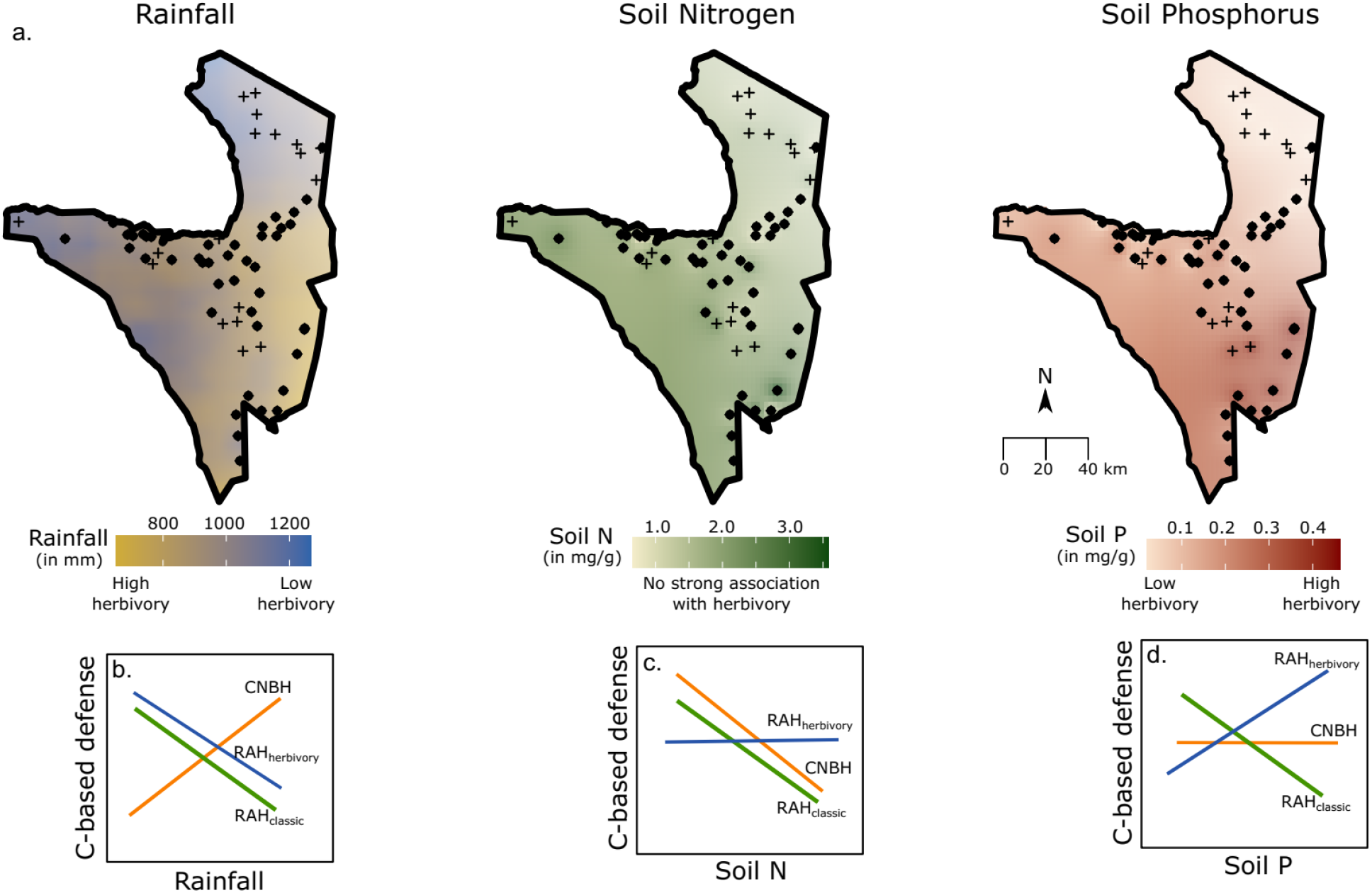
Illustration of the different resource gradients and predicted patterns in carbon-based defense traits. a) Variation in rainfall, total soil N, and total soil P across the Serengeti National Park. ‘ +’ indicate all 61 sites sampled and dots indicate sites with *Solanum incanum*. The associations of herbivory intensity with the different resources are based on previous research from the park (Mohanbabu and Ritchie 2022). The predictions from Carbon Nutrient Balance hypothesis (CNBH), classic Resource Availability Hypothesis (RAH_classic_) and herbivory dependent RAH (RAH_herbivory_) for C-based defenses along b) rainfall; c) total Soil N; and d) total Soil P gradients.

Even though the other plant defense hypotheses treat resources generically, they may offer a broad framework for predicting resource-defense relationships. The Resource Availability Hypothesis (RAH) (Coley et al. 1985, Endara and Coley 2011) posits that the species adapted to low resource environments are slow growing and hence invest heavily in defenses to reduce tissue loss to herbivores, whereas species in high resource environments are fast growing and potentially compensate for herbivory. Similarly, individuals of a widely distributed species may be adapted to different resource environments and therefore, can show well-defended phenotypes at low resource sites and poorly defended phenotypes at high resources (classic RAH applied to intraspecific variation; RAH_classic_) (Fig 1, green line). In a recent review, Hahn and Maron 2016 explicitly predict that such a negative association between resource and defense traits within species should be more likely for species adapted to physiologically stressful conditions (e.g., species found only in low resource environment). While both these predictions stem from an assumption of constant herbivory across resource gradients, researchers have shown that plants growing in high resource environments may experience greater rates of herbivory (Mattson 1980, Oksanen et al. 1981, Moreira et al. 2015, La Pierre and Smith 2016). This bottom up control of herbivory by resources may actually promote well-defended plants at high resource environments due to heightened risk of herbivory (henceforth, herbivory dependent RAH; RAH_herbivory_) (Hahn and Maron 2016, Koffel et al. 2018, Hahn et al. 2018)(Fig 1 blue line).

But increasing resources may not always increase herbivore abundance/intensity, especially if herbivores are more sensitive to other factors such as temperature or predation risk (Ritchie 2000, Anderson et al. 2010, Veldhuis et al. 2020), thereby indirectly impacting resource-defense relationships. For example, increasing precipitation in grasslands and savannas is known to increase plant growth but reduce mammalian herbivory intensity (Borer et al. 2020, Mohanbabu and Ritchie 2022) (Fig 1). Whether this mismatch in the impact of a resource on plant growth and herbivory risk results in increasing defense traits along a rainfall gradient due to the ‘excess’ availability of C (CNBH) or decreasing defense traits due reduced benefits of defense under low herbivory remains poorly understood. Furthermore, the resource-herbivory associations may also depend on the size of the herbivores (e.g., insect vs small or large mammal)(Olff et al. 2002) and their feeding habits (specialist vs generalist)(Lankau 2007, Ali and Agrawal 2012) thus highlighting the need to study patterns in defense traits in systems dominated by different types of herbivores and along multiple resource gradients.

These hypotheses are less explored in systems dominated by large mammalian herbivores (Bryant et al. 1983, Edenius 1993, Gowda et al. 2003), and plant defense response to mammalian herbivory may differ from that of the more heavily researched response to insect herbivory (Carmona et al. 2011). Firstly, chemical defenses may be less effective against mammalian herbivores that may mix secondary metabolites from multiple consumed plant species and thus dilute the impact of any single chemical (Mattson 1980). Secondly, mammalian consumption of whole or large portions of an individual plant in a single bite poses a different set of challenges to plants than invertebrate herbivory (Sanson 2006) and possibly results in different types of anti-herbivore strategies, such as avoidance through divaricate branching or structural defenses such as spines (Hanley et al. 2007, Tomlinson et al. 2016, Coverdale et al. 2019, Wigley et al. 2019). Unlike most chemical defenses, resources allocated to physical defenses cannot be metabolically reverted to resources for future plant growth and reproduction. Thirdly, the resources, predation risk and environmental conditions (e.g., temperature) that determine insect herbivore abundances (or herbivory intensity) and defenses may be different than those of mammalian herbivores. These factors inherently affect the cost-benefit ratio of allocation to defenses and consequently may influence the “dominant” type of defense depending on the identities of herbivores (Perkovich and Ward 2022) and resources but have rarely been tested for ecosystems with mammalian herbivory.

Mammalian herbivory risk can also be harder to estimate given their mobile nature and movement over larger spatial scales. This can be further complicated in natural ecosystems that have migratory herbivores and/or a diverse assemblage of herbivores, as a single timepoint measure of abundance of herbivores or damage to plants may not provide an accurate representation of the risk experienced by plants over the entire growth period. Additionally, expression of plant defense traits may lag in response to herbivory experienced during previous growing seasons (Young et al. 2003). Therefore, we propose an alternative method for estimating herbivory risk using satellite-based estimates of standing biomass that allow us to study the influence of herbivory intensity on plant traits across an entire landscape.

In this paper, we studied intraspecific variation in three types of C-based defenses: spine density, phenolic content, and lignin content, at 43 sites for a widely distributed perennial herbaceous plant species, *Solanum incanum*. These sites spanned substantial natural gradients of rainfall (a proxy for water availability) and total soil N and P (Ruess and Seagle 1994, Anderson et al. 2007) in the Serengeti National Park, Tanzania. Additionally, as these resources may be associated with mammalian herbivore abundance and herbivory intensity, we also estimated ecosystem-level large herbivore herbivory intensity (HI) at these sites. We then assessed associations between the different plant defense traits, plant resources and HI and compared our results with the predictions from different plant-defense theories for each of the resources (Fig. 1). We specifically addressed whether a) herbivory intensity varies with browser and grazer abundance, and along multiple resource gradients, b) defense traits vary along gradients of herbivory intensity and one or more plant resources, and c) resources and herbivory intensity have independent associations with defense traits.

## Methods

Field site: The Serengeti National Park in northern Tanzania is among the last remaining contiguous grassland savanna ecosystems in the world. It is home to a highly diverse group of resident and migratory large mammalian herbivores including wildebeest (*Connochaetes taurinus*), plains zebra (*Equus quagga*), Thomson’s and Grant’s gazelles (*Eudorcas thomsonii, Nanger soemmerringií*), impala (*Aepyceros melampus*), eland (*Taurotragus oryx*), elephants (*Loxodonta africana*), topi (*Damaliscus lunatus*) and Coke’s hartebeest (*Alcelaphus buselaphus*) (Sinclair and Norton-Griffiths 1979, McNaughton 1985). The plants in East Africa have co-evolved with these large herbivores for at least 3 million years (Sinclair et al. 2008) and potentially have adaptations that help them survive intense herbivory pressures with losses often ranging between 60-90% of aboveground biomass (McNaughton 1985).

Study organism: *Solanum incanum* (hereafter *Solanum*), is a pan-African and pan-Asian herbaceous plant which is consumed by several browser and mixed-feeding herbivore species (hereafter “browsers”) such as impala, elephant, gazelles and eland (Kartzinel et al. 2015, Coverdale et al. 2019, Veldhuis et al, unpubl.). Even within the Serengeti, *Solanum* is widely distributed and is almost as common as *Digitaria macroblephara* and *Themeda triandra*, two of the most abundant grasses in the park. Unlike the grasses, *Solanum incanum* invests in both physical and chemical defenses. Their stems have spines which can vary drastically in density and have been shown to be inducible by herbivory (Coverdale et al. 2019). Additionally, it produces both phenolics and alkaloids, the latter of which is mostly present in the fruits. In terms of the leaf structural content, like other plants, they also invest in structural carbon such as lignin and cellulose which may also confer some anti-herbivore properties. The wide geographic distribution, in addition to allocating to different types of defense traits makes *Solanum incanum* a ‘model’ species to explore intraspecific spatial variation in response to abiotic and biotic factors.

Study Design: From 2000 to 2002, several sites (n=102) across the national park were identified to form a network of vegetation survey plots-the trans-Serengeti plots (Anderson et al. 2007). These plots encompass much of the substantial spatial variation in rainfall (700-1125mm), total soil N (0.2-3.7 mg g^-1^) and total soil P (0.007-0.53 mg g^-1^) across Serengeti National Park (Ruess and Seagle 1994, Anderson et al. 2007).

Plots were sampled by randomly dropping points in 10 X 10 km grids and choosing one accessible point per grid such that the plot network includes a wide range of variation in abiotic factors. In 2018, we sampled 61 of these plots of which *Solanum* was present in 43 of them (Fig 1a), only slightly less than the abundant grasses, *Digitaria macroblephara* (at 50 sites) and *Themeda triandra* (at 47 sites). At each site, we sampled five individuals of *Solanum* that were at least 5m apart. For each individual, we recorded spine density as the number of spines on a 5cm length of stem at approximately 5cm from the base and we also collected 5 fully expanded mature leaves which were dried and shipped to Syracuse University, USA for further analyses. We collected soil cores at each site to characterize soil nutrient availability. Total soil N and P were estimated at the Sokoine University of Agriculture, Tanzania using the Kjeldahl method and the persulfate digestion method, respectively. We use a ten-year average for rainfall at these sites that was extracted from the Climate Hazard Group InfraRed Precipitation with Station data (CHIRPS) database (Funk et al. 2015) which provides high resolution precipitation estimates based on both long-term climate averages and weather station data.

### Estimating Herbivory intensity (HI)

Herbivory intensity is defined as the proportion of plant biomass that is consumed by herbivores. To obtain site-specific measures, we estimated HI as 1-ratio of Vegetation Index-based standing biomass and rainfall-based measures of primary productivity.

Primary productivity: We predicted primary productivity across the park using established relationships between rainfall and herbaceous primary productivity. Briefly, we combined data on biomass production and rainfall from several grasslands globally (Gill et al. 2015, Veldhuis et al. 2016), including data from the Serengeti (Sinclair 1975, McNaughton 1985, Ritchie 2014, Veldhuis et al. 2019) and estimated the slope and intercept for the relationship between biomass productivity and growing season rainfall (Fig S1). This yielded the following equation with R^2^ =0.83:

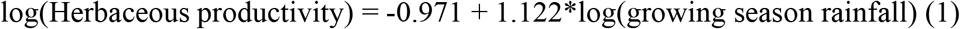

We then predicted park-wide herbaceous productivity using spatiotemporal rainfall data from the CHIRPS data (Funk et al. 2015). Growing season rainfall estimates were calculated by summing monthly CHIRPS rainfall estimates for the period September-May on a pixel bases with a resolution of 5×5 km using linear models (lm()). Primary productivity was then predicted for each year using equation 1.

Standing biomass: To estimate end of season herbaceous biomass at the park-scale we used Enhanced Vegetation Index (EVI) provided by MODerate-resolution Imaging Spectroradiometer (MODIS) with an approximate 250 × 250 m pixel resolution and 16-day interval between 2009 and 2018 (MOD13Q1) (Didan 2015). We chose the Julian dates (129,145 and 161) as they correspond to the end of the season (Fig S2). We corrected for signals from non-herbaceous evergreen species and soil differences by subtracting the minimum value of EVI (in a year) from all the EVI estimates. To confirm the usefulness of EVI estimates in predicting plant biomass in the Serengeti region, we regressed remotely sensed EVI values against ground-measured biomass data from grazed plots at 8 exclosure sites from 4 non-consecutive years (N=32).

We found that EVI Julian day 145 corrected for minimum EVI proved to best explain the variation in standing biomass (Table S1) and was therefore used for subsequent analyses. End-of-season biomass was then calculated for each year at a resolution of 250×250m at the park-scale using the following equation (2) obtained from the regressions (R^2^=0.45) described above:

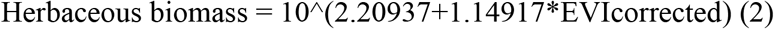

Park-wide 10-year average herbivory intensities from 2009-2018 were then calculated using the following equation:

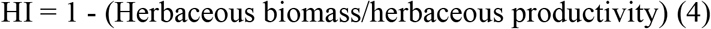

We then extracted values of HI for each of the 61 sampled sites. All spatial analyses were run in R3.5.1 (R Core Team 2018) using the raster package (Hijmans et al. 2021).

Herbivory Intensity (HI) is essentially an ecosystem-level estimate of the impact of multiple herbivore species on herbaceous biomass and should correlate with herbivore abundance (Staver et al. 2021). As data on herbivore abundance are unavailable for our sites of interest, we tested for associations between HI and herbivore abundance over a narrower region in the central Serengeti, for which multi-year camera trap data are available through the Snapshot Serengeti project (Anderson et al. 2010, Swanson et al. 2015). Briefly, we gathered count data for different herbivore species from the 225 camera traps deployed between 2010 to 2013 and distributed over a 1125km^2^ region. Using the “consensus_data.csv” file, we summed over all browser and mixed-feeding species known to consume *Solanum* species and divided it by the number of days the cameras were deployed, to account for varying search effort among cameras. As there is an inherent scale mismatch between the field of view of the camera and the resolution (250m × 250 m) of the satellite-based herbivory intensity (HI), and errors that inevitably result from estimating animal abundances from aggregated individuals (Burton et al. 2015, Pacifici et al. 2019), we averaged browser abundances over multiple cameras within 5 × 5 km grid cells. To match scales of herbivore abundance and HI, we averaged the HI estimates within 5 × 5 km grids. We then compared the average herbivore abundance with satellite-based estimates of HI for *N* = 58 grid cells. We repeated the analysis for predominantly grazing species (wildebeest, zebra, buffalo, topi, hartebeest, and gazelles) to test if HI was better associated with risk to graminoids. Given the scarcity of data on large herbivore distribution at the landscape scale, we argue that EVI-based estimate of herbivory intensity provides a proxy for risk from herbivory at different sites.

### Chemical defense traits

We estimated total foliar phenolic content using the Folin-Ciocalteu assay (Ainsworth and Gillespie 2007). Briefly, 5mg of the dried and ground leaf sample per individual was homogenized in 2ml of ice-cold 95% v/v Methanol and centrifuged at 13000g for 5min at room temperature after 48 hours of incubation in the dark. 100ul of the supernatant was diluted with 100ul of distilled water and 200ul of 10% (v/v) Folin-Ciocalteu reagent (Sigma Aldrich) was added and mixed thoroughly. After 2 minutes, 800ul of 700mM sodium carbonate solution was added to each tube and the reaction mixture was incubated at room temperature for 20 minutes. One ml of the reaction mixture was transferred to a cuvette and absorbance at 765nm was recorded for blanks, gallic acid standards and samples. Phenolic content was calculated as gallic acid equivalents based on the standard curve. Although Solanum may also produce alkaloids, they disintegrate quickly unless stored at −80 C, which limited our ability to quantify alkaloids.

We estimated structural lignin content of *Solanum* leaves using a multi-step sequential digestion with ANKOM 200 Fiber Analyzer (ANKOM Technology, New York USA). The leaf samples for each site were pooled and ~0.5g of the dried leaf material was sealed in a special 25u porous bags from ANKOM. The bags were weighed and sequentially treated with neutral detergent solution at 100 °C for 75 minutes, acid detergent solution at 100 °C for 60 minutes and 98% sulfuric acid at room temperature for 3 hours. Between each step, the bags were washed with hot water and acetone to remove any remaining detergents, dried at 105 °C overnight and weighed. Finally, the filter bags were ashed at 500 °C for 330 minutes in a muffle furnace. The difference in the weights between conc. sulfuric acid treatment and ashing were used to estimate the percentage of lignin per dry mass of plant tissue.

### Statistical Analysis

All the data analyses were run in R (R Core Team 2020).

Association between herbivory intensity (HI) and herbivore abundance: We ran a linear (lm()) with HI as the dependent variable and log transformed herbivore abundance as the independent variable. We also included a quadratic term for herbivore abundance to account for potential non-linear associations.

#### Variation in HI

To understand variation in herbivory intensity in response to different resource gradients we used data from all 61 sites that were sampled. We ran a linear model with HI as the dependent variable and rainfall, total soil N and P along with their two-way interaction terms as the independent variables. As some of the resource gradients in the Serengeti are moderately correlated (Pearson correlation coefficients: rainfall-soil P= −0.48, soil N-soil P=0.62, rain-soil N= −0.06), we also analyzed the data using univariate models to specifically understand relationships between herbivory intensity and each resource gradient.

#### Variation in defense traits

As sites (n=43) are the experimental units in this study, we averaged data for the 5 individuals per site for all traits except lignin, for which samples were pooled at the site level before ANKOM analyses. To understand associations of defense traits with herbivory intensity (HI) we used linear models, lm(), between a given defense trait and herbivory intensity. Additionally, to explore variation in defense traits along gradients of resource availability, we ran linear models using lm() with a given defense trait as the dependent variable and rainfall, total soil N and P as the independent variables plus their two-way interactions. Finally, to parse out the association of resources with defense traits, that is independent of their indirect association with HI, we added HI and its interaction with resources to the previous model but dropped previously non-significant interaction terms to avoid over-parameterizing models. If the associations between resource and defense traits are no longer found to be significant after HI and its significant interaction terms are included, the results suggest an indirect link between resources and defense traits mediated by an association of HI with plant resources. If the defense-resource associations continue to be significant, it would suggest that resources may influence plant defense traits independently of HI, perhaps through mechanisms highlighted in the RAH and CNBH hypotheses.

For the purposes of visualizing associations of defense traits with different resources, we present partial residual plots (Fig 3) of variation in both dependent (log(spine density+1)) and independent (rainfall, total soil N and soil P) variables once the covariance between the remaining independent variables is accounted for.

**Figure 2:**
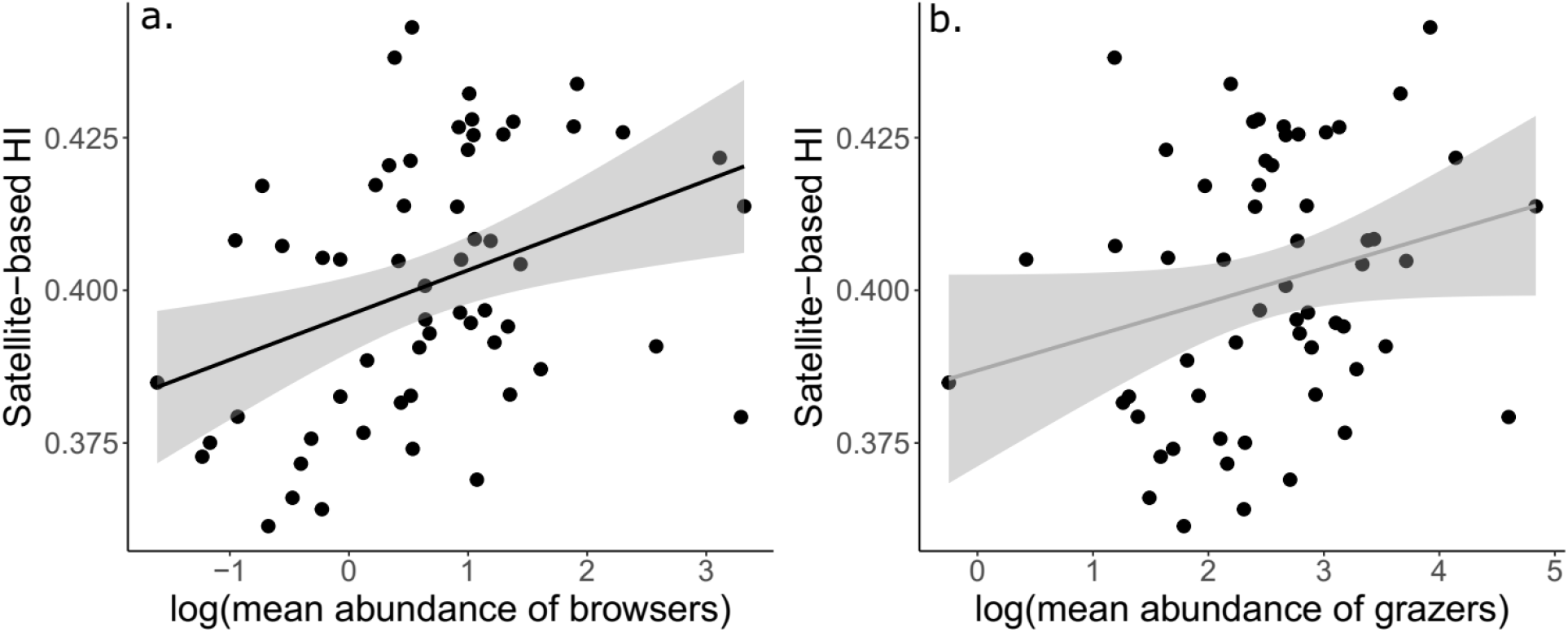
Association between satellite-based herbivory intensity (HI) and log-transformed a) browser abundance (species that consume *Solanum*), and b) grazer and mixed-feeding species abundance based on the Snapshot Serengeti camera trap data. The dots represent abundance and HI estimates in 5 × 5 km in the central Serengeti, dark and grey lines represent significant and insignificant relationships, respectively, and grey areas represent the 95% confidence intervals.

**Figure 3:**
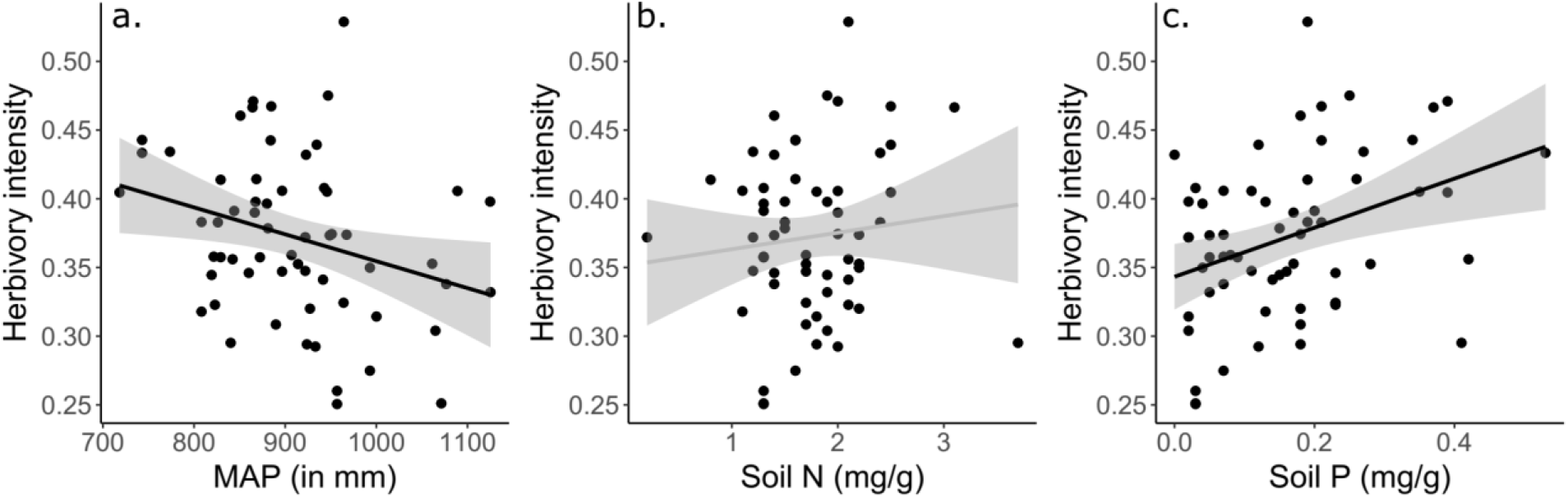
Variation in herbivory intensity along gradients on a) Mean annual precipitation (in mm), b) Total soil N (%), and c) Total soil P (%). In the figure, each dot is the estimated herbivory intensity at a site, dark lines represent significant relationships and grey lines represent insignificant relationships based on univariate models, and grey areas represent the 95% confidence intervals.

## Results

Herbivory intensity varied from 0.25 to 0.53, implying that 25 to 53% of the aboveground biomass was consumed by herbivores across the 43 sites at which *S. incanum* was present. Although the regions with camera traps spanned a narrower range of HI from 0.36 to 0.44, HI was positively associated with browser and mixed-feeding herbivore abundances (slope (SE): linear term: 0.012 (0.004), p=0.001; insignificant quadratic term: −0.003 (0.002), p=0.08, R^2^=0.15) (Fig 2a), but not for grazer and mixed-feeding herbivore abundances (0.006 (0.003), p=0.06, R^2^=0.04) (Fig 2b).

HI was negatively associated with rainfall (−0.18(0.08), p=0.02) (Fig 3a), positively associated with soil P (0.02(0.007), p=0.003) (Fig 3c) and not associated with soil N (0.006 (0.007), p=0.40) (Fig 3b) in univariate models. These landscape patterns are also corroborated by results from a long-term grazing exclosure experiment in the Serengeti (Mohanbabu and Ritchie 2022). Including all the resources in a single model however, we found that soil P was the only variable to show an independent, though only marginally significant, trend with herbivory intensity (0.022 (0.012), p=0.06) (Supplementary info Table S2).

All three defense traits showed substantial intraspecific variation: spine density ranged from no spines (undefended) to ~72 spines per 5 cm, and total phenolic and structural lignin content varied by about a factor of three, ranging between 1.75-8.77 nmol of gallic acid equivalents/mg of sample, and 4.57-13.43%, respectively. Spine density was bimodally associated with herbivory intensity such that well-defended plants were present at both high and low HI (HI^2^= 7.30 (1.53), p<0.001; HI=-6.98(1.53), p<0.001) (Fig 4a). In contrast, variation in total phenolic content (−0.38(0.28), p=0.19) (Fig 4b) and lignin (0.54(0.34), p=0.12) (Fig 4c) was not associated with variation in herbivory intensity.

**Figure 4:**
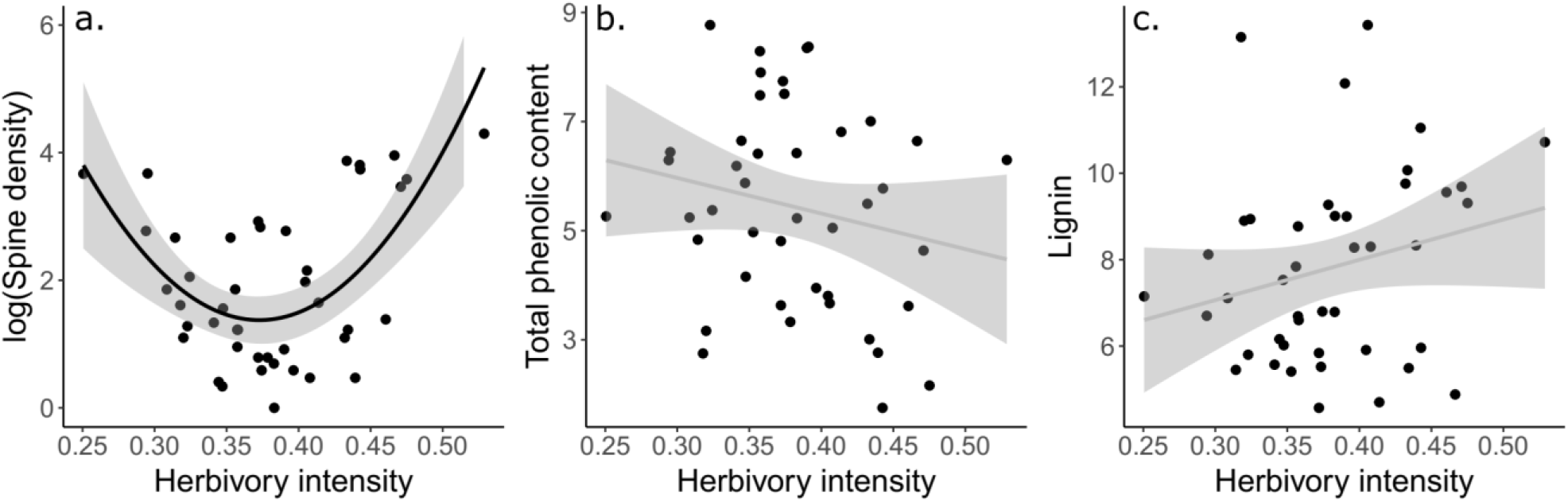
Variation in defense traits a) Spine density (number of spines/ 5cm of stem), b) Total phenolic content (nmol of gallic acid equivalent/ mg of plant dry mass), and c) Lignin content (in % of plant dry mass) with herbivory intensity measured from Enhanced Vegetation Index. In the figure, each dot is a site, dark lines represent significant relationships and grey lines represent insignificant relationships based on univariate models, and grey areas represent the 95% confidence intervals.

Resources also explained some of the variation in defense traits. Spine density increased with both rainfall (0.48 (0.21), p=0.03) (Fig 5a, Table 1a) and soil P (0.59(0.28), p=0.04) (Fig 5c, Table 1a), but did not change with soil N (−0.08 (0.24), p=0.73) (Fig 5b, Table 1a). Total phenolic content was negatively associated with rainfall (−0.67(0.37), p=0.08) (Table 1a), although the association was only marginally significant. Variation in soil nutrients did not contribute to explaining variation in total phenolic content. Lastly, lignin did not vary along gradients of resources (Table 1a). Finally, to test for direct effects of resources on defenses, as opposed indirect effects via HI, we included HI in the resource-defense models. For all three defenses, including herbivory intensity in the same model as resources renders the resource-defense associations insignificant, except for spine density and soil P, which remains positive and significant (Table 1b).

**Figure 5:**
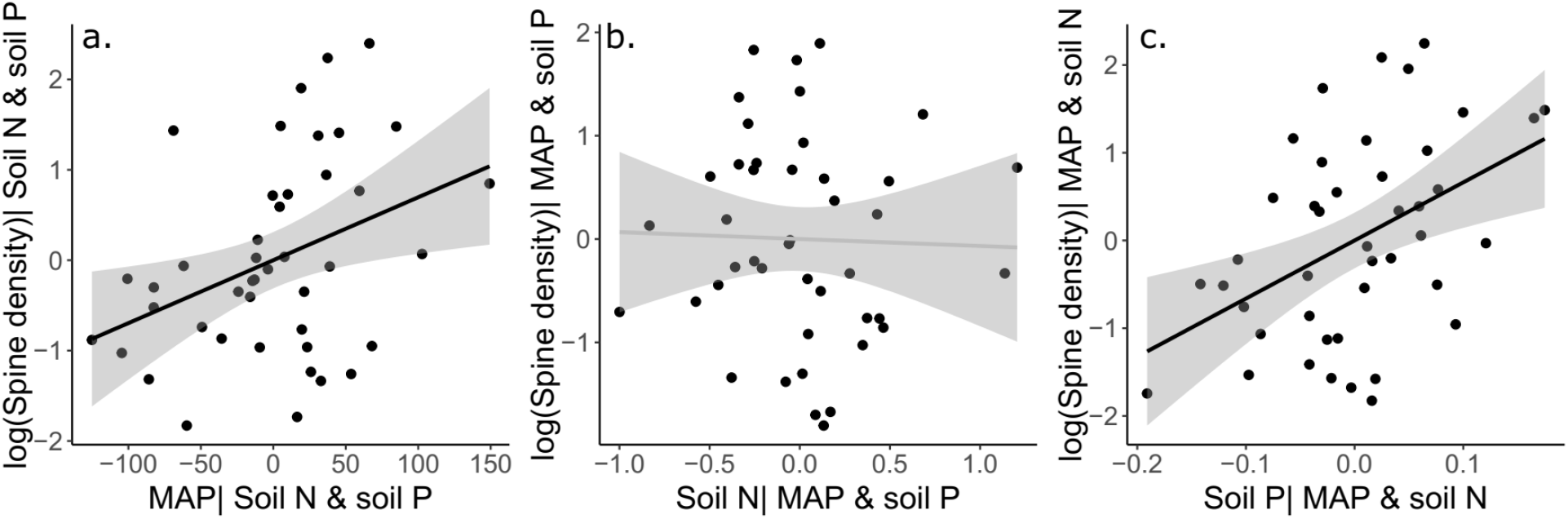
Variation in spine density along gradients of a) rainfall, b) soil N, and c) soil P represented as partial residual graphs, i.e. covariation due to other variables in the model are already accounted for (see Methods). In the figure, each dot is the log of mean spine density at a site, dark lines represent significant relationships and grey lines represent insignificant relationships based on univariate models, and grey areas represent the 95% confidence intervals.

**Table 1:** Summary of the linear models for each of the defense traits spine density, total phenolic content and lignin for a) Resource model and b) Resource and Herbivory intensity (HI) model. For the models with both resources and HI, all the interaction terms between resources and resource and HI were insignificant and were dropped to simplify the model. Bold values indicate p < 0.05 and italics represents 0.1>p>0.05

| Defense trait | Factors | Spine density | Total phenolic<br>content | Lignin |
| --- | --- | --- | --- | --- |
| a) Resource models | MAP | **0.48(0.21)** | *-0.67(0.37)* | 0.14(0.46) |
|  | Soil N | -0.08(0.24) | 0.45(0.42) | -0.50(0.52) |
|  | Soil P | **0.59(0.28)** | -0.56(0.49) | 0.95(0.61) |
|  | MAP x Soil N | 0.22(0.30) | 0.43(0.53) | -0.42(0.66) |
|  | MAP x Soil P | -0.29(0.19) | 0.25(0.33) | 0.58(0.41) |
|  | Soil N x Soil P | 0.23(0.18) | 0.11(0.31) | -0.15(0.39) |
| b) Resource and HI | MAP | 0.24(0.18) | -0.47(0.35) | 0.09(0.43) |
|  | Soil N | -0.13(0.20) | 0.29(0.40) | -0.27(0.49) |
|  | Soil P | **0.61(0.23)** | -0.45(0.47) | 0.34(0.58) |
|  | HI | **-6.19(1.54)** | -0.32(0.31) | 0.45(0.38) |
|  | HI2 | **6.34(1.54)** |  |  |

## Discussion

We explored patterns of intraspecific variation in defense traits along multiple resource and herbivory gradients in a plant species that can experience substantial herbivory by large mammals. We found that spine density was positively associated with both rainfall and total soil P in the system. These positive resource-defense associations suggest potential for resources to influence herbivores and their associated risk of plant damage, thereby supporting the RAHherbivory hypothesis (Hahn and Maron 2016, Koffel et al. 2018), and result in well-defended plants at high resource supply. To our knowledge, this is the first-time total soil P has been shown to be associated with a physical defense trait even though P is not a major structural component of largely carbon-based defense traits. Our results for phenolics and lignin contrast with most prior research, which generally shows positive associations between climate factors or N supply and chemical defenses (Moreira et al. 2015, Hahn and Maron 2016, Abdala-Roberts et al. 2016, Hahn et al. 2018). Thus, our work highlights the potential for varied responses of defense traits to three common resources that limit plant growth, which may have greater influence on plant-herbivore interactions in the future, given projected climate change and anthropogenic inputs of nutrients (Peñuelas et al. 2013, IPCC 2014).

Interestingly, the positive association between soil P and spine density is only partially explained by the indirect influence of resources mediated by positive associations between herbivory intensity and soil P. The RAH_herbivory_ hypothesis posits that plant defenses should increase with resource supply, perhaps due to increased plant growth at resource-rich sites which may support higher herbivore biomass and herbivory intensity (Hahn and Maron 2016). Although soil P and herbivory intensity (HI) are positively associated as expected (Fig 3c), soil P continues to explain variation in spine density even when HI is included in linear models. Thus, P may influence allocation to defense through other mechanisms. A probable alternate mechanism could be elevated levels of photosynthesis and C assimilation at high P conditions, which is then allocated towards a C-based physical defense, such as spine density. However, the lack of associations of the other two C-based defenses with soil P (Table 1) is not consistent with this alternative hypothesis. More work is needed to elucidate mechanisms by which P may affect defense allocation.

In contrast to soil P, increasing spine density at higher rainfall (Fig 5a) is consistent with the Carbon-Nutrient Balance Hypothesis (CNBH). In grasslands like the Serengeti, rainfall, productivity and plant C:N ratios are strongly associated (Sinclair 1975, McNaughton 1985). Increasing rainfall allows for higher rates of CO_2_ uptake relative to N- or P-based biochemical machinery. Therefore, plants may be nutrient limited rather than C-limited at high rainfall sites, allowing C to be allocated to C-based structural defense at relatively lower opportunity costs (Bryant et al. 1983, Hamilton et al. 2001). Alternatively, the positive association between rainfall and spine density may also occur if high herbivory intensity is supported at high rainfall sites. However, satellite-based estimates of HI are negatively related to rainfall in univariate models. This observation also contradicts the assumptions of most resource-defense models (Loreau and Mazancourt 1999, Hahn and Maron 2016, Koffel et al. 2018). It is, however, possible that the relatively nutrient rich *Solanum* may experience higher herbivory risk compared to the neighboring background of low-quality grasses, thereby resulting in well-defended *Solanum* individuals.

Surprisingly, spine density increased at both low and high herbivory intensity. This pattern may be a by-product of the negative association of soil P (and herbivory intensity) with rainfall in our study system. As a result, low HI sites are also associated with higher rainfall, and within-plant availability of C. This C might be allocated to defenses despite relatively low herbivory rates. In contrast, high HI at high soil P and low rainfall and C availability sites may select for better defended plants to deter herbivores even if allocating C for defense may have higher costs. Therefore, defenses maybe favored either where resources for defense are abundant under low but still present risk or where resource-expensive plant tissue is at high risk. This hints at the possibility of a shift from a resource-associated (bottom-up) to an herbivory-associated (top-down) control of allocation to defenses along an environmental gradient.

Effects of large mammalian herbivores on plant defenses have been harder to address at the landscape scale due to the difficulties associated with measuring risk from herbivores. Although we were able to show that herbivory intensity was positively associated with browser relative abundance, the relationship was weak, as might be expected. Camera traps often feature high sampling variance for mobile, highly aggregated animals, and detection probabilities may vary with species identity, herd size or animal behavior (Treves et al 2010, Burton et al 2015, Kolowski and Forrester 2017). Moreover, the central Serengeti where the camera traps are placed is a region with relatively high herbivore activity and had a narrow range of HI (0.36 to 0.45) as compared to the range in the rest of our study (0.25 to 0.55). This coupled with the fact that estimation of HI contains errors from the relationship between rainfall and ungrazed biomass driven in part by litter accumulation may also have limited our ability to find a stronger correlation. Yet, we detected statistically significant associations between HI and browser abundance, and between HI and spine density in Solanum, which suggests that HI is an important axis of variation for plant defenses, likely due to its correlation with browser abundances. Additionally, the lack of association between HI and grazer abundance also suggests that HI is a reliable measure of risk from browsing rather than grazing species, likely because EVI-based estimates of plant biomass may be disproportionately affected by herbaceous vegetation which tends to be greener than the surrounding grasses. Thus, vegetation index-based HI offered us a chance to explore patterns in defense traits along a gradient of risk from herbivores that was independent of the plant defense levels, and provides an alternative method for estimating herbivory intensity.

The lack of variation in lignin and phenolics with resource supplies or herbivory intensity may indicate that those traits may not be as strongly influenced by mammalian herbivory as are spines. Firstly, both phenolics and lignin may be responding to insect herbivory or microbial pathogens, rather than mammalian herbivory. Data from other savannas (Davies et al. 2016) suggests that invertebrate herbivores contribute approximately 10-20% to overall herbivory and these numbers are likely to be even smaller in the Serengeti where herbivory by large mammals is exceptionally high (McNaughton 1985). Secondly, both phenolics and lignin are leaf-level carbon-based defenses and may get allocated to labile (total phenolics) or structural (lignin) carbon based on other factors such as variability in herbivory or leaf phenology (e.g., leaf lifespan) (Coley et al. 1985). Despite both defense traits exhibiting no association with plant resources, phenolics were indeed negatively associated with lignin (Fig S3), which suggests a potential tradeoff in allocating carbon to these different compounds. Finally, the two defenses may contribute to plant fitness in ways not necessarily related to defense, such as when phenolics protect against light damage (Isah 2019, Erb and Kliebenstein 2020) and lignin provides structural support and extends leaf lifespan (Kitajima et al. 2012).

Our work adds to a limited number of field studies on the responses of defenses to multiple resources in the presence of mammalian herbivores. Our field survey limits our power of inference and field experiments that manipulate different resource levels and herbivory are likely necessary to determine cause-effect relationships of resources and herbivory risk on defense traits. Although our analysis is limited to a single species, *S. incanum* represents a type of “model” plant for intraspecific variation studies because it exhibits multiple types of defenses, is common in an unusually wide variety of environments (i.e., was present at 43 of the 61 sites) and is consumed by multiple mammalian herbivore species (Kartzinel et al. 2015, Coverdale et al. 2019, Veldhuis et al, unpubl.). Even though our work maybe easily extended to a vast majority of species in Solanaceae that invest in different types of defenses, future work on different plant families will help evaluate any generality of our results. Finally, it is possible that the herbivores and defense traits are responding to the supply of a different resource such as Na or Ca, but across site comparison of these elements in the soil show low correlation with that of soil P in the Serengeti. Regardless, variation in defense traits along gradients of other resources such as Na and Ca are still unclear and a worthy topic for future research.

In conclusion, we show substantial intraspecific variation in defense traits of *Solanum incanum*, especially spine density, that is associated with both variation in herbivory intensity and environmental resources. Interestingly, spine density was positively associated with soil P even after accounting for herbivory intensity, which emphasizes the influence of resources other than C and N on plant defenses. In contrast, a three-fold variation in phenolic and lignin content was unexplained by variation in resources or HI. Our work indicates that future research should consider both different types of defenses and resources in order to better understand the complex link between resource supply and allocation to defenses.

