## Supplementary information for "Landscape variation in defense traits along gradients of multiple resources and mammalian herbivory"

Table S1: Results from linear regressions between log-transformed plant biomass from the LTGE experiment and corrected (suffix '.cor') or uncorrected Enhanced Vegetation Indices (EVI) for different Julian days (129,145,161).

| Y variable | X variable | Intercept | Estimate (SE) | Mult R2 | Adj R2 | F | df |
|---|---|---|---|---|---|---|---|
| log10 (biomass) | EVI.145.cor | 2.209 | 1.149 (0.222) | 0.47 | 0.45 | 26.8 | 1,30 |
| log10 (biomass) | EVI.161.cor | 2.228 | 1.523 (0.318) | 0.44 | 0.42 | 22.8 | 1,28 |
| log10 (biomass) | EVI.129.cor | 2.199 | 1.062 (0.240) | 0.39 | 0.37 | 19.5 | 1,30 |
| log10 (biomass) | EVI.161 | 2.078 | 1.402 (0.304) | 0.43 | 0.41 | 21.2 | 1,28 |
| log10 (biomass) | EVI.145 | 2.093 | 1.096 (0.221) | 0.45 | 0.43 | 24.6 | 1,30 |
| log10 (biomass) | EVI.129 | 2.105 | 0.979 (0.236) | 0.36 | 0.34 | 17.1 | 1,30 |

Note: All associations were statistically significant at alpha=0.05. EVI.145.cor had the highest Adjusted R2 and was therefore used in further analyses.

Table S2: Results from a multivariate model between herbivory intensity and multiple resources and their interactions. This analysis was performed on all 61 site irrespective of the presence of Solanum at those sites.

| Factor | Estimate (SE) | P value |
|---|---:|---:|
| Intercept | 0.37(0.009) | <0.001 |
| MAP | -0.007(0.010) | 0.48 |
| Soil N | 0.001(0.010) | 0.91 |
| *Soil P* | *0.022(0.012)* | *0.06* |
| MAP * Soil N | 0.018(0.014) | 0.20 |
| MAP * Soil P | -0.001(0.008) | 0.86 |
| Soil N * Soil P | -0.0001(0.0082) | 0.99 |

### The adjusted $R^2$ for the full model was 0.11 with p=0.054

Figures

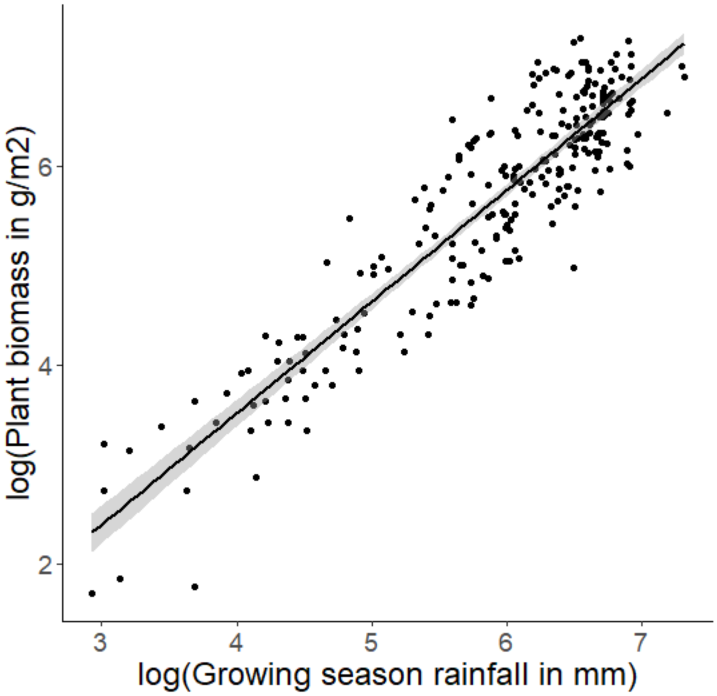


Figure S1: Relationship between rainfall and plant biomass in fenced plots from various sites across the globe ($R^2$=0.83).

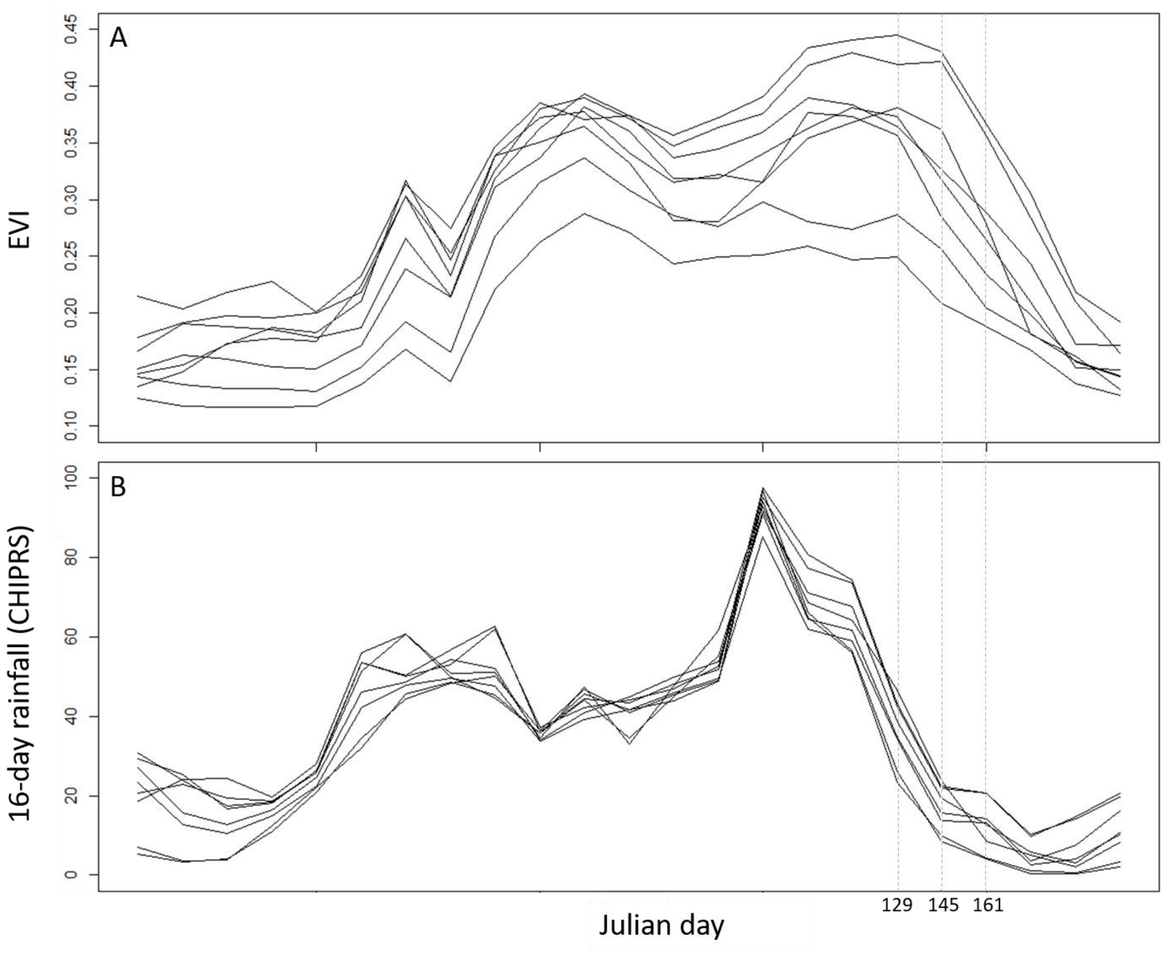


Figure S2: Mean annual variation in EVI (A) and 16-day rainfall (B) for the 8 Long Term Grazing Exclosure sites between 2000-2016. In May (Julian days 129 and 145), rainfall declines marking the beginning of the dry season followed by a drop in EVI and NDVI in June (Julian day 161).

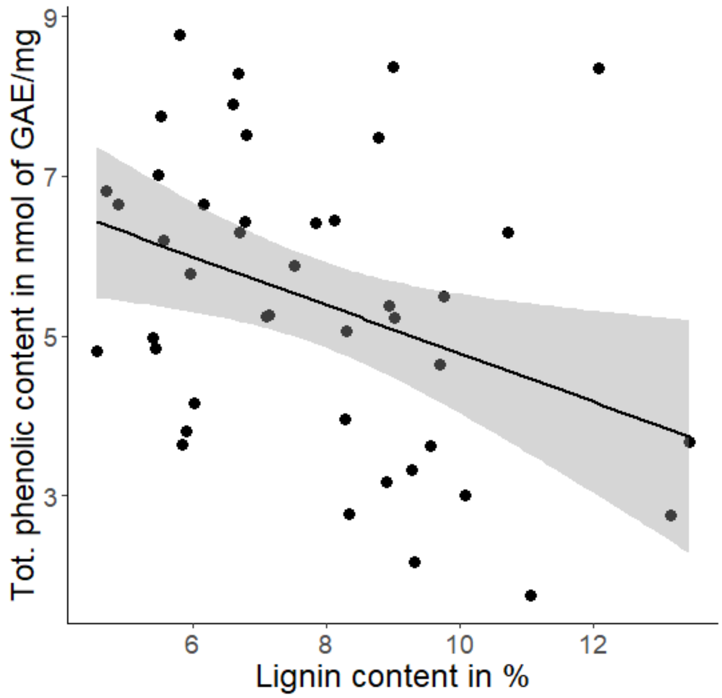


Figure S3: Trade-off between Lignin and phenolics in *Solanum incanum*. The Pearson correlation coefficient is -0.37 (p=0.02). Dots represent different sampling sites and shaded area represents 95% CI.
